# Neurodevelopmental Impact of Bipolar Disorder Genetic Risk on Cortical Thickness and Network Topology in Adolescents

**DOI:** 10.1101/2025.06.07.658415

**Authors:** Xiaobo Liu, Lang Liu, Jiadong Yan, Jinzhi He, Bin Wan, Ruiyang Ge, Jinming Xiao, Guoyuan Yang

## Abstract

Bipolar disorder (BD) is a highly heritable psychiatric condition characterized by recurrent mood episodes that commonly manifest during adolescence. Although polygenic risk scores (PRS) effectively quantify genetic susceptibility to BD, the neurobiological correlates of this genetic risk during adolescent brain development remain unclear. Using normative modeling and longitudinal neuroimaging data from the Adolescent Brain Cognitive Development (ABCD) cohort (N = 4519), we examined cortical thickness (CT) deviations and structural covariance network (SCN) alterations in adolescents stratified into high and low BD genetic risk groups based on PRS. Adolescents with high PRS for bipolar disorder exhibited greater cortical thickening in the inferior frontal gyrus and primary visual cortex, whereas those with low PRS demonstrated greater thickening in the posterior cingulate cortex and middle precentral gyrus. These patterns suggest PRS-related variations in cortical maturation, potentially reflecting distinct neurodevelopmental trajectories associated with genetic susceptibility to bipolar disorder. Furthermore, high PRS individuals displayed altered SCN topology, characterized by decreased local clustering and enhanced global network efficiency. Longitudinal data show that these abnormal regions exhibit atypical developmental trajectories, accompanied by a global reorganization of network topology. Additionally, high genetic risk was associated with lifestyle factors, especially correlated with increased positive expectancies toward substance use. Eventually, we found three potential BD risk genes during adolescence, including PLEKHA2, ZSCAN31 and ANK3, encoding the function of the postsynaptic membrane and synaptic membrane. These findings elucidate early neurodevelopmental deviations linked to genetic risk for BD, highlighting potential biomarkers for early identification and targeted interventions.

## Introduction

Bipolar disorder (BD) is one of the most heritable mental illnesses characterized by recurrent episodes of mania or hypomania and depression, often manifesting during adolescence (Gerard & Buehler, 2004; Gordovez & McMahon, 2020; MacQueen et al., 2005). Adolescents with a high genetic predisposition to BD frequently exhibit subclinical symptoms and cognitive impairments prior to clinical onset, making this population crucial for investigating early neurodevelopmental alterations (Hafeman et al., 2017; Smoller, 2020). Recently, polygenic risk scores (PRS) have been increasingly utilized in psychiatric genetics research (Choi et al., 2020; Lewis & Vassos, 2020), which derived from genome-wide association studies (GWAS), quantifying an individual’s genetic susceptibility to complex disorders (Uffelmann et al., 2021). Previous studies have demonstrated that individuals with high genetic risk for severe mental disorders may already exhibit impairments in impulse control and heightened sensitivity of the reward system prior to meeting full clinical diagnostic criteria (Alloza et al., 2020; Whalley et al., 2012, 2013). In BD, PRS can aggregate the cumulative effects of multiple genetic variants to predict individual disease risk (Mullins et al., 2021), which was associated with widespread cortical morphological abnormalities (Cattarinussi et al., 2022). These neurobehavioral alterations may serve as critical contributing factors to the increased engagement in maladaptive behaviors during adolescence, such as cigarette smoking, alcohol consumption, and cannabis use. However, despite extensive research on BD, the neuroimaging impact of genetic susceptibility on brain development in large-scale developmental cohorts remains poorly understood.

Neuroimaging studies have consistently documented both localized anatomical changes and widespread network-level dysfunction in BD (van Erp et al., 2018; Xia et al., 2022). Regionally, reductions in cortical thickness—particularly within the medial prefrontal cortex and anterior cingulate cortex—have been frequently observed, implicating circuits involved in affect regulation and executive function (Rimol et al., 2010; Xia et al., 2022). In addition, fronto-limbic circuit dysfunction has been implicated in impaired emotion regulation, while heightened responsivity in fronto-striatal circuits is thought to underlie aberrant reward sensitivity (Alamian et al., 2017; Frangou, 2012). A typical functional connectivity extends to sensory and affective processing regions, potentially contributing to destabilized emotional, cognitive, and motor control (Perry et al., 2019). While resting-state functional connectivity has provided valuable insights into the large-scale network alterations in BD, its test-retest reliability, susceptibility to transient brain states, and limited structural interpretability pose challenges for understanding stable, trait-like neurodevelopmental disruptions (Syan et al., 2018). In this context, structural covariance analysis offers a stable and complementary approach that reflects inter-regional coordination of brain morphology across individuals, partially recapitulates functional networks, and is thought to capture long-term developmental and genetic influences on brain organization (Alexander-Bloch et al., 2013; Kim et al., 2020; Zielinski et al., 2010). Therefore, assessing structural covariance in relation to polygenic risk provides a promising avenue to characterize macroscale network alterations linked to genetic vulnerability for BD.

Traditional case-control studies often rely on group-level statistical comparisons, which may overlook individual variability and fail to capture the heterogeneous neurobiological profiles, particularly in the context of BD. In contrast to the traditional case-control framework, normative modeling provides a robust approach for capturing individual-level deviations by referencing a normative population, thereby enabling the investigation of neurobiological heterogeneity at the single-subject level (Rutherford et al., 2022). The strength of normative modeling approach lies in its ability to make statistical inferences at the level of the individual, which may enable accurately personalized assessments of neurobiological abnormalities in BD. Until recently, the normative modeling has been successfully applied to various neuropsychiatric conditions, revealing individualized (Ge et al., 2024a; Lv et al., 2021) deviations that better encapsulate the biological heterogeneity inherent in psychiatric disorders (Lv et al., 2021a; Verdi et al., 2023). However, there remains a lack of studies examining normative model deviations in brain structural morphology in BD, which could provide novel insights into the highly heterogeneous neurodevelopmental abnormalities associated with the BD.

In this study, we utilized the Adolescent Brain Cognitive Development (ABCD) dataset to investigate cortical thickness deviations in adolescents at high and low PRS groups for BD using normative modeling. We further analyzed longitudinal trajectories of cortical thickness to examine developmental differences between high and low PRS groups. Additionally, we assessed structural covariance networks to explore network-level alterations associated with genetic risk. Then, we examined behavioral differences between high and low PRS groups to evaluate potential associations between neurobiological markers and lifestyle factors. Finally, we further delineated specific BD–associated genes related to cortical structural abnormalities. By integrating genetic risk stratification, neuroimaging biomarkers, and behavioral assessments, this study aims to delineate early neurobiological alterations associated with BD genetic risk and establish a framework for identifying individuals at elevated risk for developing BD. These findings have the potential to inform early identification strategies and contribute to the development of precision psychiatry approaches for BD.

## Results

### PRS distribution and clinical symptom differences

To investigate the neurodevelopmental impact of genetic susceptibility to BD, we defined high and low PRS groups from a sample of 4519 participants ( aged 119.37 ± 7.34 months, 2117 females/2402 males). The PRS was computed using a weighted sum of risk alleles associated with BD in genome-wide association studies (GWAS), following standard methodology (Choi et al., 2020). We used a normal distribution of PRS values from the study population and applied predefined percentile-based cutoffs, including 5% and 95% (Taylor et al., 2019; Whalley et al., 2013), to classify individuals into two groups : high PRS (*N* = 237, aged 118.27 ± 7.20 months, 101 females /136 males) and low PRS (*N* = 237, 120.54 ± 6.97 months, 114 females/123 males) (**Fig. 1a**).

**Figure 1.**
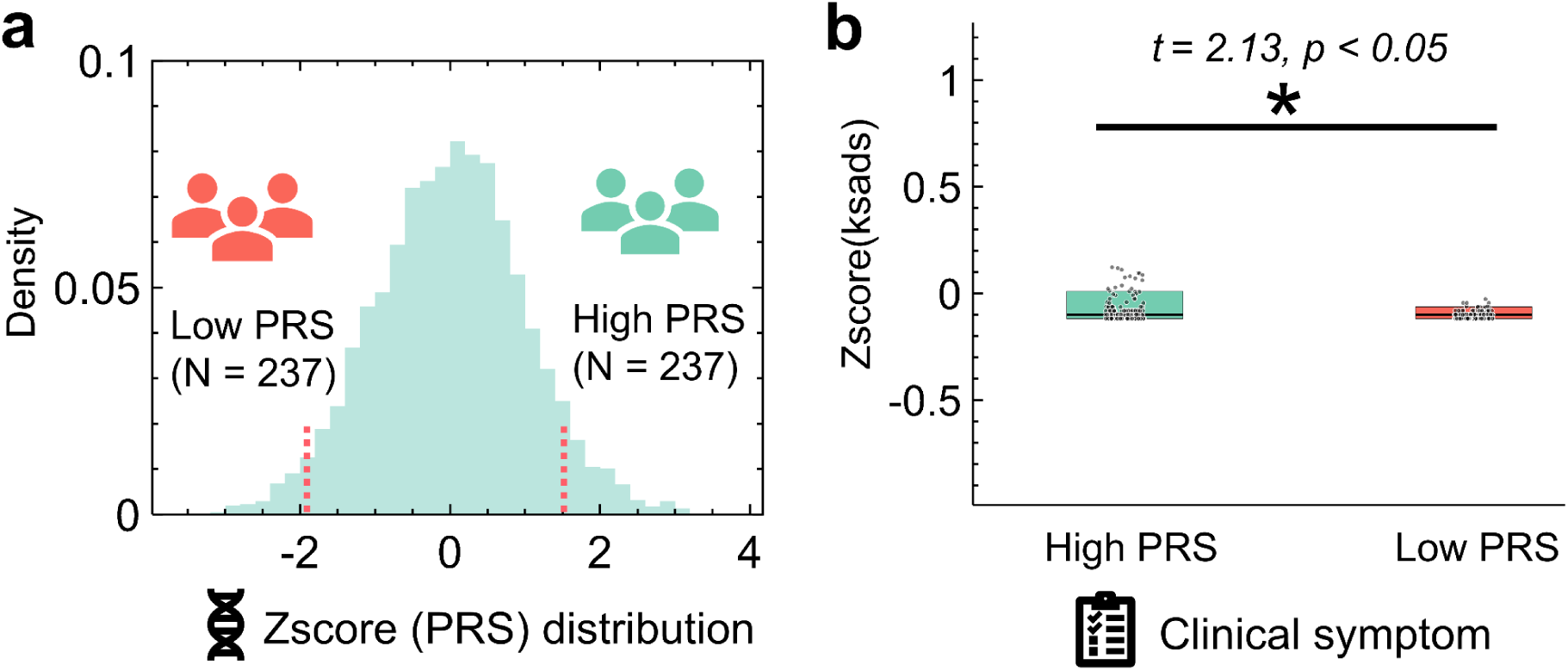
The adolescence with bipolar disorder (BD) risking based on Polygenic Risk Score (PRS) and Clinical Symptoms. (**a**) PRS distribution across the study population. The histogram represents the density distribution of PRS values, with participants divided into two groups: low PRS (N = 237, red) and high PRS (N = 237, green), based on the indicated thresholds (Participants falling within the highest 5% and lowest 5% of the PRS distribution). (**b**) Comparison of clinical symptom severity (Z-score of *KSADS* assessment) between high and low PRS groups. The boxplot illustrates group differences, with individual data points overlaid. A significant difference is observed between the high and low PRS groups (*t* = 2.13, *p* < 0.05). The asterisks indicate statistically significant differences (*p* < 0.05).

To assess whether genetic risk influences subclinical psychiatric symptoms, we compared the Z-scored KSADS (Kiddie Schedule for Affective Disorders and Schizophrenia) between the high and low PRS groups. The KSADS assessment provides standardized clinical measures of mood disorder symptoms, and Z-scores were computed to normalize symptom severity across participants. A two-sample *t*-test revealed a significant difference in clinical symptom severity between the groups (*t =* 2.13, *p <* 0.05) (**Fig. 1b**). Specifically, adolescents in the high PRS group exhibited elevated subclinical symptom severity compared to those in the low PRS group, suggesting a potential link between genetic risk and early psychiatric manifestations.

### Heterogeneous cortical thickness deviations in high and low BD risking populations

To further investigate the neurobiological impact of genetic risk for BD, we applied normative modeling to quantify individual-level deviations in cortical thickness between high and low PRS groups. This approach allows for a more personalized assessment of neurodevelopmental alterations compared to traditional case-control comparisons, which often fail to capture the heterogeneity within psychiatric populations (Marquand et al., 2016).

We first computed cortical thickness deviations for each participant using a normative model trained on a large-scale reference population (Ge et al., 2024a). Normative models of cortical thickness were developed using multivariate fractional polynomial regression (MFPR) on data from 37,407 healthy individuals (aged 3–90 years) across 87 datasets spanning Europe, Australia, the USA, South Africa, and East Asia (Ge et al., 2024a). This model provided individual Z-scores representing deviations from the expected cortical thickness patterns. We then compared these deviations between the high PRS and low PRS groups, identifying regional patterns of cortical thickening and thinning (**Fig. 2a**).

**Figure 2.**
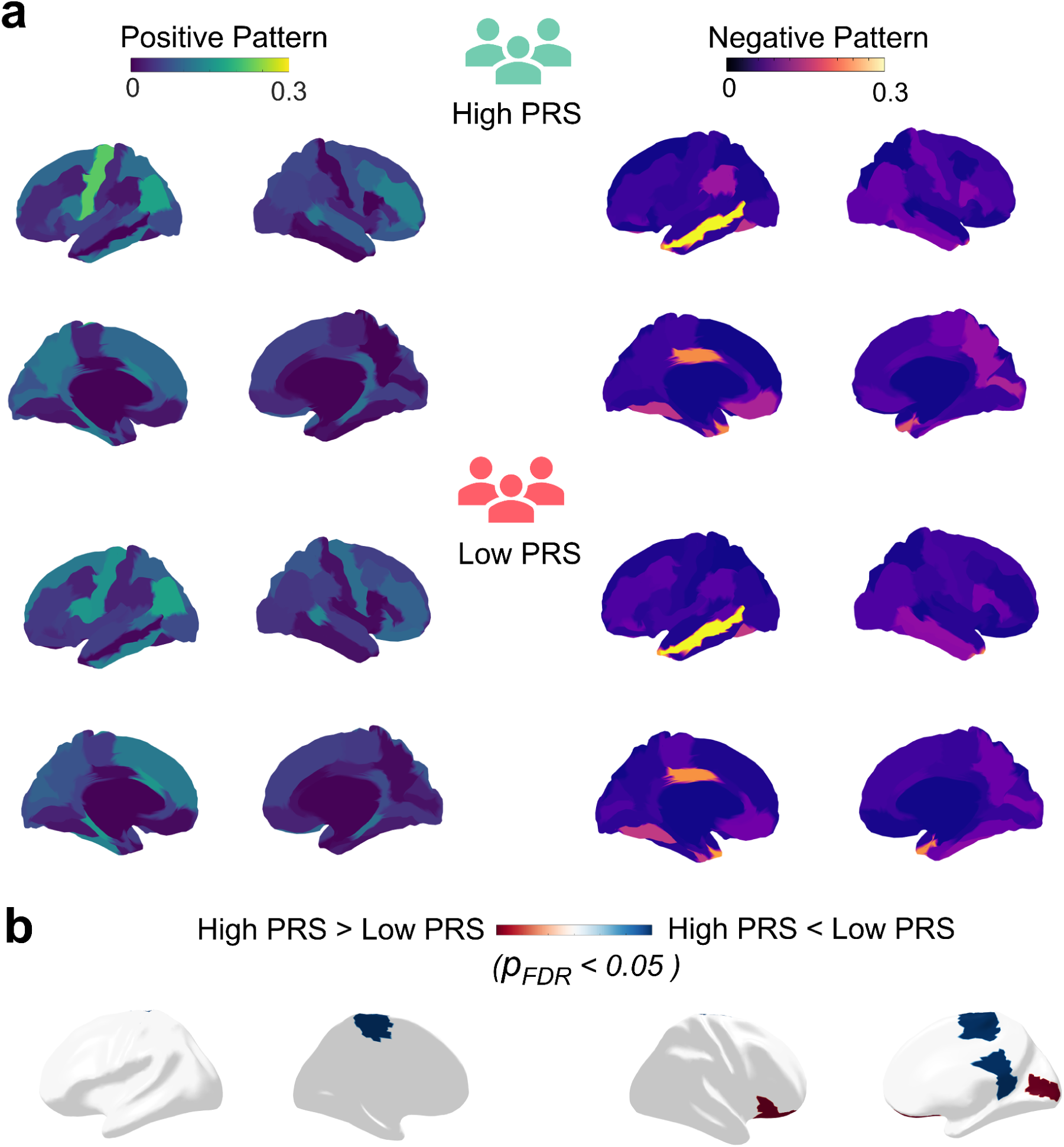
Normative model based cortical thickness deviation between high and low PRS groups. (**a**) Cortical thickness patterns illustrate differences between high PRS individuals (green) and low PRS individuals (red). The left panel shows positive pattern (ratio of participants whose cortical thickness deviation was larger than 1.96) in yellow-orange, while the right panel depicts negative pattern (ratio of participants whose cortical thickness deviation was less than -1.96) vin blue-purple. In the high PRS group, significant cortical thickening is observed in the dorsolateral prefrontal cortex (DLPFC), parietal cortex, and superior frontal gyrus (SFG), whereas significant cortical thinning is evident in the temporal pole, orbitofrontal cortex (OFC), and anterior cingulate cortex (ACC). (**b**) Direct comparison of cortical thickness deviations between high and low PRS groups. The color bar represents group differences, where red indicates greater deviations in the high PRS group compared to the low PRS group, while blue denotes greater deviations in the low PRS group. Notably, individuals with a high PRS demonstrated significantly greater cortical thickening in the inferior frontal gyrus (IFG) and primary visual region, whereas those with a low PRS exhibited increased thickening in the posterior cingulate cortex (PCC) and middle precentral gyrus (*p_FDR_* < 0.05). These differences highlight distinct neurodevelopmental trajectories between risk groups.

In the high PRS group, deviated cortical thickening was observed in the dorsolateral prefrontal cortex (DLPFC), parietal cortex, and superior frontal gyrus (SFG), whereas cortical thinning was most pronounced in the middle temporal gyrus (MTG), orbitofrontal cortex (OFC), and anterior cingulate cortex (ACC). The low PRS group also exhibited regional deviations from normative cortical thickness trajectories. Positive deviations were observed in brain regions that partially overlapped with those identified in the high PRS group, such as the dorsolateral DLPFC and SFG. Similarly, negative deviations were evident in the temporal pole, OFC, and ACC, showing a high degree of spatial concordance with the high PRS group. These shared anatomical regions suggest that, despite differences in the magnitude of deviation, both high and low PRS groups may converge along similar neurodevelopmental axes, potentially reflecting a common vulnerability architecture modulated by genetic load.

To directly compare cortical thickness deviations between the high and low PRS groups, we computed group-level differences (**Fig. 2b**). The high PRS group showed greater thickening in the inferior frontal gyrus (IFG) and and primary visual region, while the low PRS group exhibited greater thickening in the posterior cingulate cortex (PCC) and middle precentral gyrus (*p_FDR_ <* 0.05). These findings indicate group-specific cortical maturation differences, potentially linked to neurodevelopmental processes influenced by genetic risk for BD.

### Structural covariance network topography in high and low PRS populations

To further examine network-level alterations associated with genetic risk for BD, we analyzed the structural covariance network topology in high and low PRS groups. Structural covariance networks reflect the interregional correlations in cortical morphology and provide insights into brain-wide organizational differences driven by genetic susceptibility (Alexander-Bloch et al., 2013). We constructed individual structural covariance network based on cortical thickness covariance and examined two key graph-theoretic properties: path length, which measures the average shortest connection between brain regions and reflects global network efficiency, and clustering coefficient, which quantifies the degree of local interconnectedness and reflects regional network segregation (Rubinov & Sporns, 2010).

We categorized the SCNs into three canonical network configurations (**Fig. 3**, top panel). Regular networks exhibit high clustering coefficients and long path lengths, representing strong local connectivity but inefficient global integration. Small-world networks maintain a balance between local clustering and global efficiency, supporting optimized information processing. In contrast, randomized networks have low clustering coefficients and short path lengths, indicating disrupted local connectivity but increased global integration.

**Figure 3.**
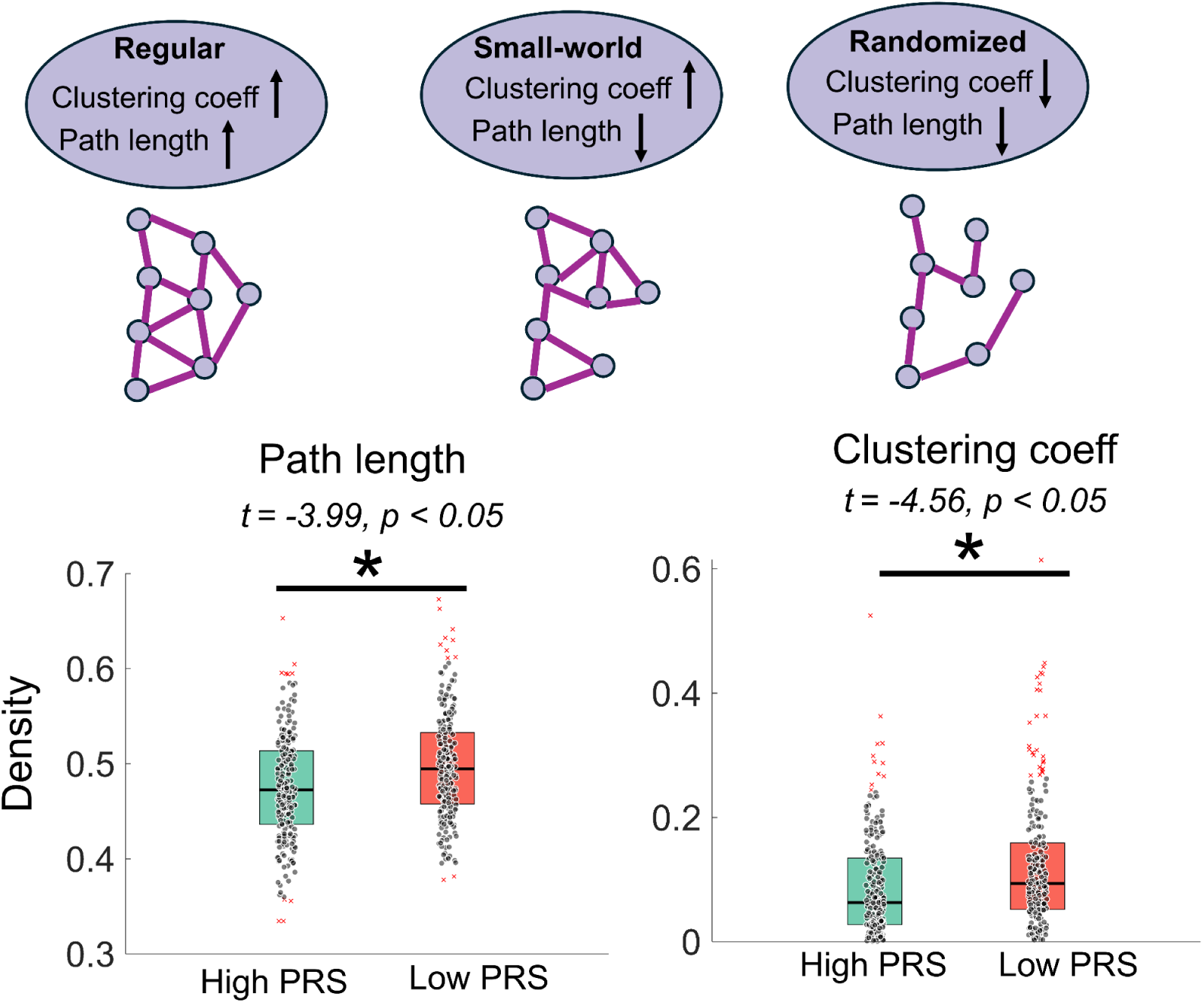
Structural covariance network topography between high and low PRS groups. This figure compares path length and clustering coefficient between individuals with high PRS and low PRS groups. The top panel illustrates different network configurations: Regular networks have high clustering coefficients and long path lengths; Small-world networks have a balance of clustering and path efficiency; Randomized networks have low clustering coefficients and short path lengths. The bottom left plot shows that the path length is significantly longer in the low PRS group compared to the high PRS group (*t =* -3.99, *p <* 0.05). The bottom right plot indicates that the clustering coefficient is also significantly higher in the low PRS group (*t =* -4.56, *p <* 0.05). The asterisks indicate statistically significant differences (*p* < 0.05).

Comparing the structural covariance metrics between groups, we found that high PRS individuals exhibited significantly shorter path lengths compared to low PRS individuals (*t =* -3.99, *p <* 0.05), suggesting that high PRS individuals exhibit more efficient global integration (**Fig. 3**, bottom left). Additionally, high PRS individuals exhibited significantly lower clustering coefficients compared to low PRS individuals (*t =* -4.56, *p <* 0.05), suggesting that high PRS individuals show reduced local network segregation (**Fig. 3**, bottom right).

### Longitudinal cortical thickness and structural network topography changes in high and low PRS groups

To further explore the developmental trajectory of cortical structure and network properties, we analyzed longitudinal changes in both cortical thickness and structural covariance network in high and low PRS groups. This approach allows us to capture how genetic risk influences cortical maturation and large-scale brain network organization over time.

First, we computed longitudinal cortical thickness changes (ΔCT) for each individual by assessing differences in cortical thickness across two time points (2-year follow up - baseline). We then compared these changes between the high PRS and low PRS groups (**Fig. 4a**). The medial central anterior gyrus (MCAG) showed significantly greater cortical thinning in the high PRS group relative to the low PRS group (*t* = 2.85, *p* < 0.05), reflecting accelerated synaptic pruning and potentially compromised maturation of regions involved in executive functioning and emotional regulation. Conversely, the superior frontal gyrus (SFG) demonstrated significantly greater cortical thickening in the high PRS group compared to the low PRS group (*t* = -2.37, *p* < 0.05), indicating reduced or atypical synaptic pruning, consistent with abnormal maturation patterns that may impact motor system development.

**Figure 4.**
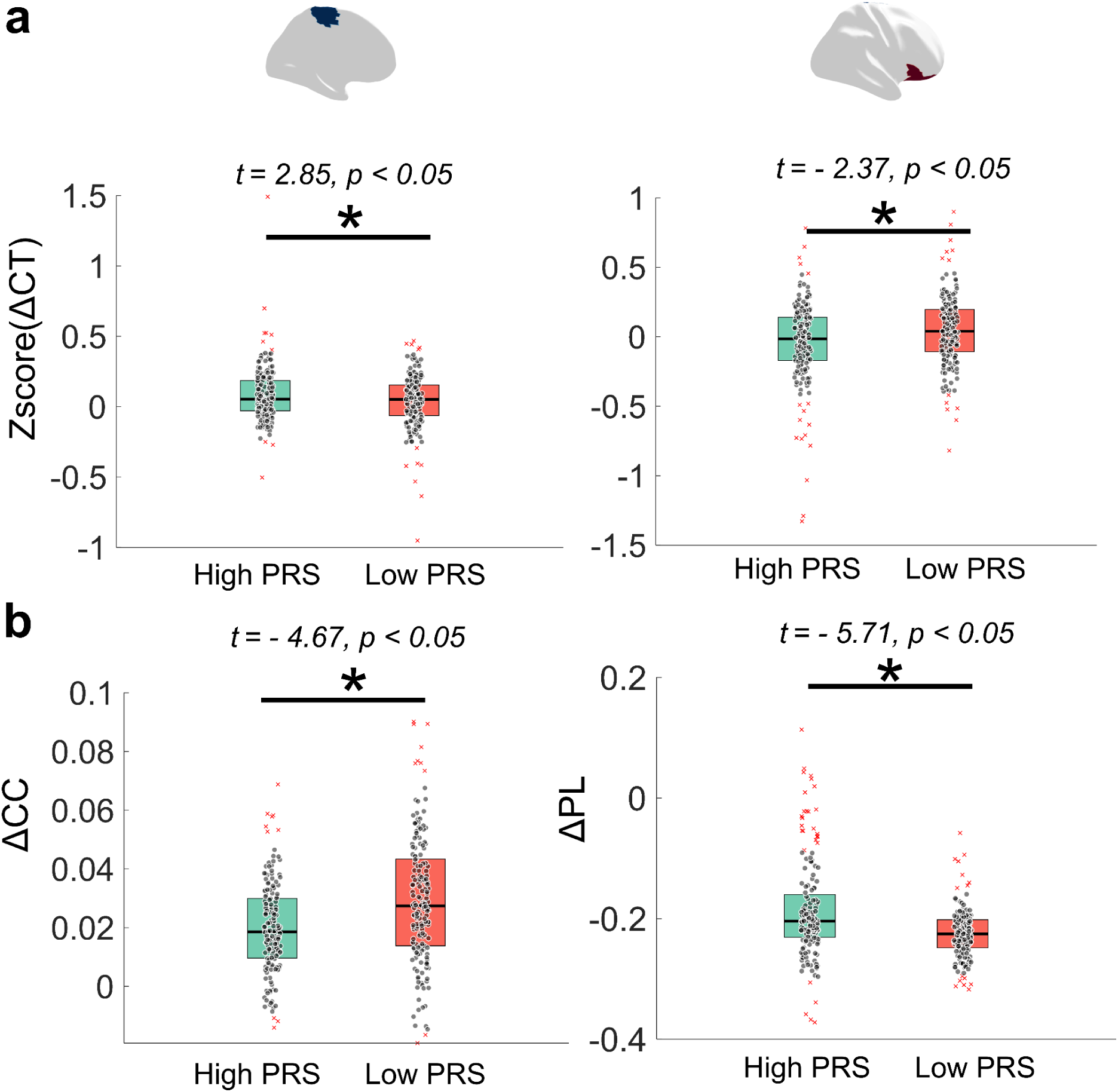
Longitudinal cortical structural changes in high and low PRS groups. (**a**) The differences in longitudinal cortical thickness changes between individuals with high and low PRS. Left panel: the middle precentral gyrus exhibits a significantly greater cortical thinning rate in the high PRS group compared to the low PRS group (*t* = 2.85, *p* < 0.05), suggesting an accelerated reduction in cortical thickness over time. Right panel: the precentral gyrus, in contrast, shows a significantly greater cortical thickening rate in the high PRS group compared to the low PRS group (*t* = -2.37, *p* < 0.05), indicating an atypical pattern of cortical development. (**b**) The longitudinal changes in structural covariance network properties between high and low PRS groups. Left panel (ΔCC): the clustering coefficient (CC), representing local network segregation, increases significantly more in the low PRS group compared to the high PRS group (*t =* 4.67, *p <* 0.05), indicating that low PRS individuals maintain stronger local connectivity over time. Right panel (ΔPL): the shortest path length (PL), which reflects global network efficiency, decreases more significantly in the high PRS group compared to the low PRS group (*t =* -5.71, *p <* 0.05), suggesting a more pronounced reorganization of global network integration in high PRS individuals. These findings suggest that genetic risk for BD is associated with distinct patterns of cortical development and network reorganization over time. The asterisks indicate statistically significant differences (*p* < 0.05).

Next, we examined longitudinal changes in structural covariance network properties by calculating differences in clustering coefficient (ΔCC) and shortest path length (ΔPL) over time (**Fig. 4b**). The clustering coefficient, which reflects local network segregation, increased significantly more in the low PRS group compared to the high PRS group (*t =* 4.67, *p <* 0.05). This suggests that low PRS individuals maintain stronger localized connectivity, which may support stable functional specialization. In contrast, the shortest path length, which measures global network efficiency, decreased significantly more in the high PRS group compared to the low PRS group (*t =* -5.71, *p <* 0.05), indicating greater reorganization of global network integration in individuals with high genetic risk.

### Association between PRS and substance use expectation

To investigate the relationship between PRS and substance use exposure, we first compared differences in positive expectancies toward alcohol and cannabis use between the high and low PRS groups. Substance use exposure scores reflect individuals’ anticipations of positive experiences related to substance use, with higher scores indicating a greater likelihood of future substance use.

Our analysis revealed significant group differences. The high PRS group exhibited significantly higher positive expectancy scores for alcohol exposure compared to the low PRS group (*t =* 2.34, *p <* 0.05; **Fig. 5a**). Similarly, positive expectancy scores for cannabis exposure were also significantly greater in the high PRS group relative to the low PRS group (*t =* 2.10, *p <* 0.05; **Fig. 5b**). These findings suggest that individuals with a higher genetic risk exhibit stronger positive expectancies toward substance use, which may contribute to an increased likelihood of future engagement in substance use behaviors.

**Figure 5.**
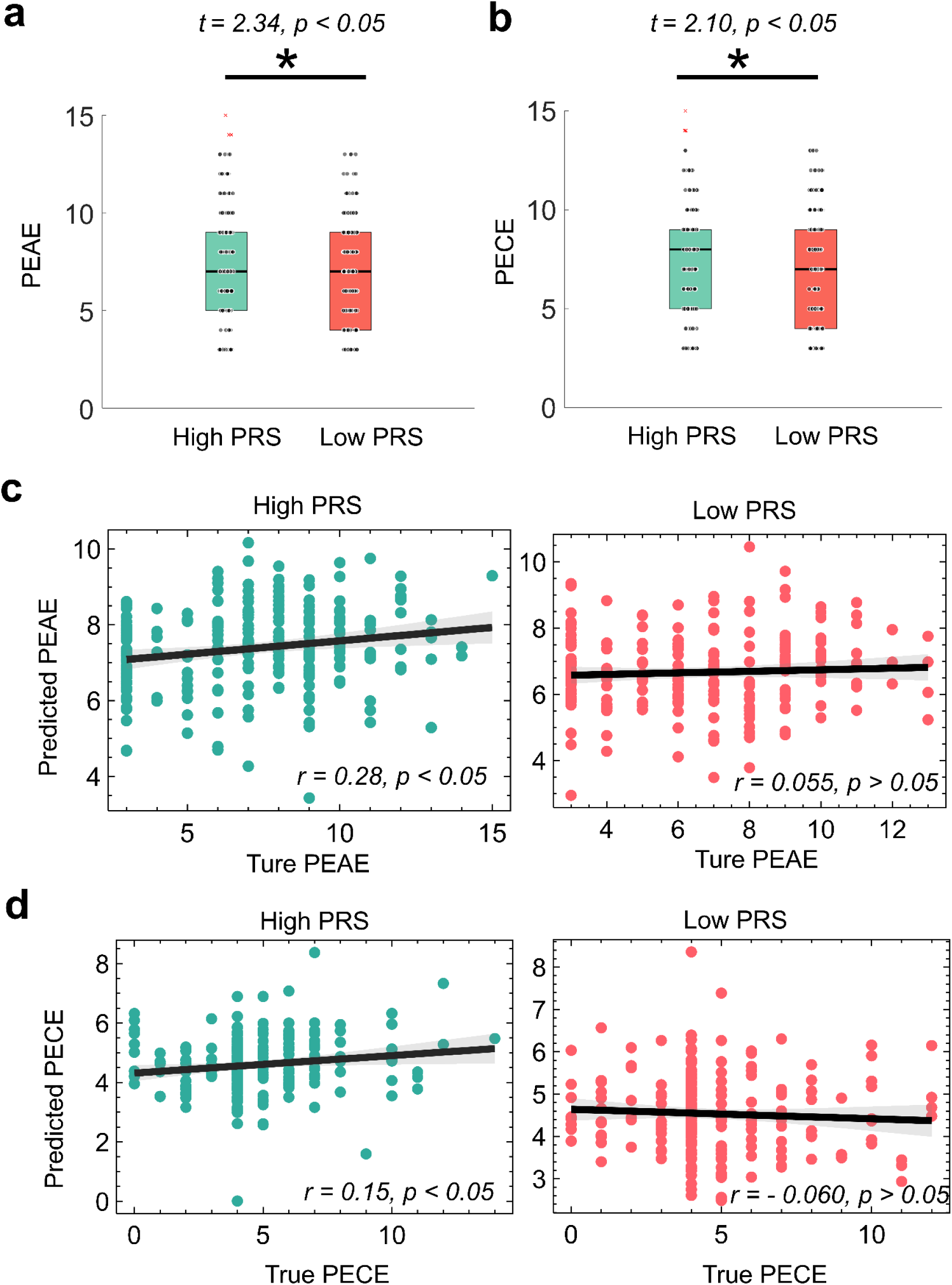
Substance use exposure between individuals with high and low PRS groups. (**a**) The differences in positive expectation for alcohol exposure (PEAE) between the high and low PRS groups, with significantly higher levels in the high PRS group (*t =* 2.34, *p <* 0.05). (**b**) The differences in positive expectation for cannabis exposure (PECE), where the high PRS group also exhibits significantly greater exposure compared to the low PRS group (*t =* 2.10, *p <* 0.05). The asterisks indicate statistically significant differences (*p* < 0.05). Boxplots illustrate the median, interquartile range, and outliers for each group. (**c**) Baseline cortical thickness in the high PRS group could predict positive expectation for alcohol exposure (*r =* 0.28, *p <* 0.05) while the low PRS group could not (*r =* 0.055, *p >* 0.05). (**d**) Baseline cortical thickness in the high PRS group could predict positive expectation for cannabis exposure (*r =* 0.15, *p <* 0.05) respectively, while the low PRS group could not ( *r =* - 0.060, *p >* 0.05).

**Figure 6.**
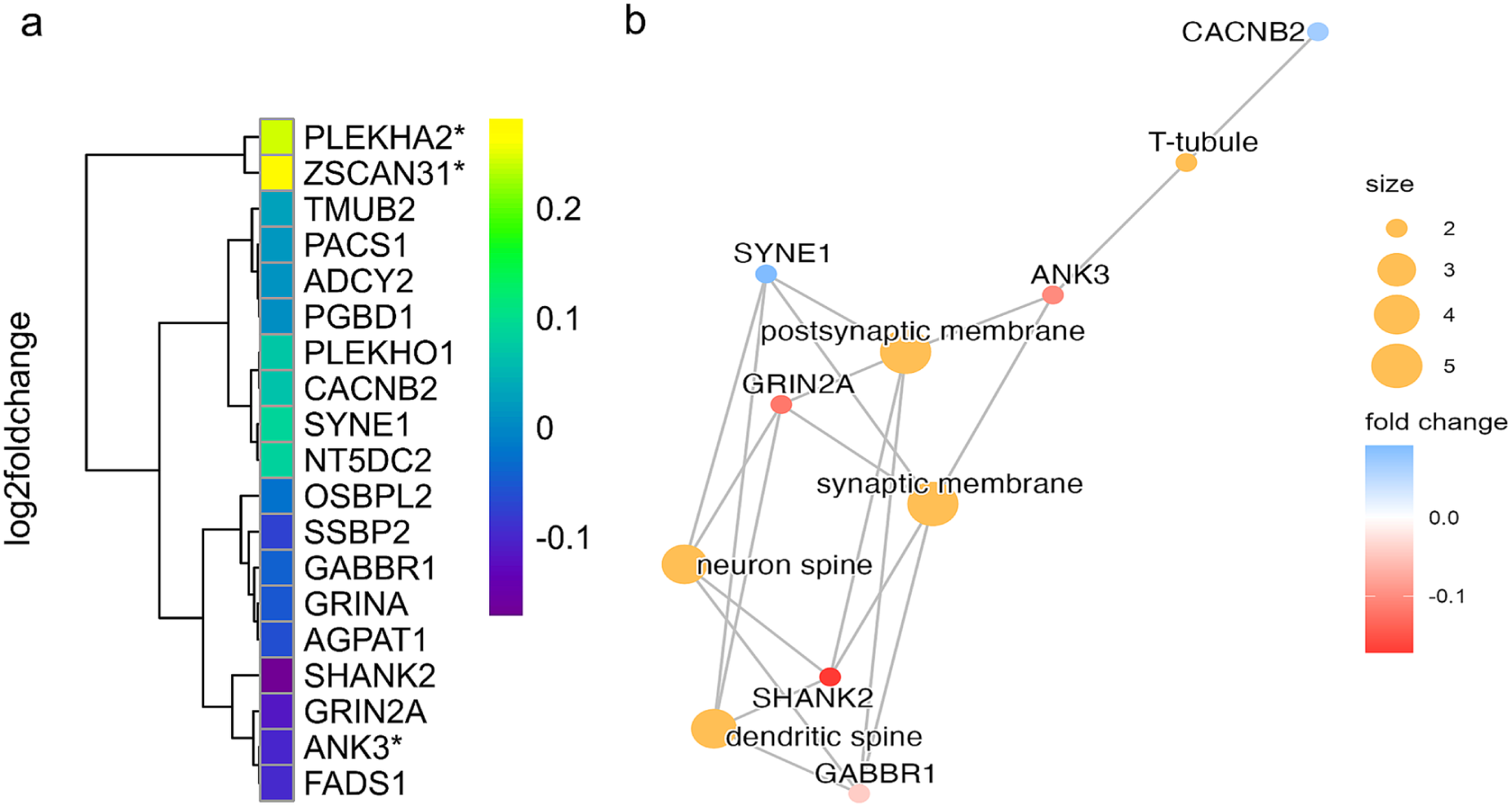
BD risk genes and pathway enrichment related to cortical thickness abnormalities. (**a**) Heatmap showing log₂ fold change of genes significantly associated with cortical thickness alterations in BD. Genes were identified by correlating regional gene expression (AHBA) with neuroimaging data and cross-referenced with BD GWAS hits. *PLEKHA2*, *ZSCAN31*, and *ANK3* (marked with asterisks) were overlapping genes between transcriptomic and genetic datasets. (**b**) Network of enriched cellular components based on spatially correlated genes using Metascape. Node size reflects term gene count; color indicates fold change. Enriched terms include postsynaptic membrane, synaptic membrane, dendritic spine, and T-tubule. *ANK3* is notably associated with synaptic structures, suggesting functional relevance to BD pathophysiology.

In addition, we employed a Support Vector Regression (SVR) model with baseline cortical thickness as the predictor to examine whether neural structural features could predict individuals’ positive expectancies for alcohol and cannabis exposure. As shown in **Fig. 5c**, in the high PRS group, the SVR-predicted scores for alcohol exposure were significantly correlated with the true exposure scores (*r =* 0.28, *p <* 0.05), whereas in the low PRS group, the association was not significant. A similar pattern emerged for cannabis exposure (**Fig. 5d**): the SVR prediction based on cortical thickness was significantly correlated with actual positive expectancy scores in the high PRS group *(r =* 0.15, *p <* 0.05) but not significant in the low PRS group. These results highlight that genetic vulnerability (high PRS) may intensify the neural substrates underlying substance use expectancies, pointing to cortical thickness as a potential biomarker for risk-related behavioral patterns.

### Genetic mechanisms in high and low PRS groups

Although GWAS have identified a set of risk alleles associated with BD (Mullins et al., 2021), these variants are often located in regulatory regions and lack functional annotation regarding their spatiotemporal expression in brain tissue. To address this limitation, we integrated regional gene expression data from the Allen Human Brain Atlas (AHBA) (Arnatkeviӗiūtė et al., 2019) and performed spatial correlation analyses between the expression patterns of BD-associated GWAS genes and cortical thickness abnormalities observed in BD patients (Markello & Misic, 2021). This approach aimed to investigate whether specific candidate genes exhibit expression patterns that are not only genetically linked to BD risk but also anatomically aligned with disease-related structural brain alterations. By doing so, we sought to enhance the mechanistic understanding of how these genes may contribute to the neurobiological underpinnings of BD. Among the BD-associated GWAS genes, we observed that multiple genes, including PLEKHA2, ZSCAN31, and ANK3, showed significant spatial correlations with the cortical thickness abnormalities identified in our imaging analyses. In addition, PLEKHA2, ZSCAN31, and ANK3 were found to be differentially expressed in bipolar disorder based on bulk RNA-seq analysis from the PsychENCODE dataset (Gandal et al., 2018; Wei et al., 2024). This suggests that the regional expression patterns of PLEKHA2, ZSCAN31, and ANK3 may contribute to the neurobiological mechanisms underlying cortical abnormalities observed in bipolar disorder. Furthermore, via pathway analysis, the most expressed genes were postsynaptic membrane, synaptic membrane, dendritic spine, neuron spine and T-tubule. ANK3 was more related to the postsynaptic membrane and synaptic membrane. These results further revealed potential BD risking genes during adolescence.

## Discussion

This study provides novel insights into the neurodevelopmental consequences of genetic susceptibility to BD during adolescence. By applying polygenic risk stratification combined with normative modeling of cortical thickness and structural covariance networks in a large developmental cohort, we identified distinct neurobiological alterations associated with BD risk. Adolescents with higher genetic susceptibility exhibited marked deviations in cortical maturation trajectories, predominantly in regions crucial for executive functioning and emotion regulation, suggesting early neurobiological vulnerability. These structural changes were accompanied by significant differences in network topology, highlighting reduced local connectivity and enhanced global integration. The presence of elevated subclinical psychiatric symptoms and increased substance use expectancies further underscores the clinical relevance of these neuroimaging markers, providing a foundation for future precision psychiatry strategies aimed at early detection and intervention.

Our imaging analyses revealed that high PRS individuals showed a pattern of cortical thinning in regions implicated in executive control and emotional regulation, while also exhibiting cortical thickening in motor-related regions like the precentral gyrus. These findings align with previous literature indicating that frontal and motor cortices are key loci for BD’s neurodevelopmental disruptions (Hibar et al., 2018; Howes & Shatalina, 2022; Maletic & Raison, 2014; Parenti et al., 2020). By highlighting regional heterogeneity in the direction of cortical deviations, our results suggest that BD genetic risk may simultaneously accelerate typical maturational thinning in certain areas and induce atypical thickening in others (Westacott & Wilkinson, 2022). This dual pattern may reflect differential synaptic pruning or compensatory neurodevelopmental processes (Petanjek et al., 2023; Tau & Peterson, 2010). Critically, these results extend prior observations by demonstrating that these structural deviations precede clinical onset in adolescents, offering an early window into BD pathogenesis and reinforcing calls for early targeted interventions.

In addition to examining regional thickness deviations, we found that high PRS individuals exhibited shorter path lengths and lower clustering coefficients in their structural covariance networks, indicative of increased global integration but reduced local segregation. This structural network-level pattern is reminiscent of randomized or less segregated network organization, echoing earlier studies linking diminished local clustering to cognitive and mood dysregulation in BD (Patankar et al., 2020; Tucker, 1991; Zhang & Raichle, 2010; Zuo et al., 2021). Our results thus suggest that BD genetic risk may drive large-scale network reorganization, shifting the brain’s balance away from stable local circuits toward a more integrated—yet potentially less specialized—configuration. This finding dovetails with previous evidence of default mode network hyperconnectivity in BD (Dobri et al., 2022; Ford et al., 2013; Kim et al., 2020; Narayanan et al., 2015), reinforcing the view that genetic predisposition can reshape global brain networks in ways that heighten vulnerability to emotion dysregulation and cognitive impairments (Dvir et al., 2014; “Emotion Processing Deficits,” 2015; Etain et al., 2008; Gräff & Mansuy, 2009; Green et al., 2007). Ultimately, such network alterations may serve as biologically informed markers for identifying high-risk individuals before pronounced clinical symptoms emerge.

Our longitudinal analyses further underscore the dynamic nature of these neurobiological changes. Specifically, high PRS adolescents demonstrated an accelerated rate of cortical thinning in frontal regions over time, accompanied by greater reorganization of network efficiency (i.e., decreasing path length). This temporal perspective aligns with reports suggesting that adolescence is a critical period during which BD-related brain deviations may become more pronounced (Liu et al., 2023; Markt et al., 2024; McWhinney et al., 2021). By tracking individuals across multiple time points, we reveal how genetic risk modulates ongoing neural development, potentially catalyzing the emergence of subclinical symptoms. These findings not only overlap with prior work documenting frontal cortex thinning in incipient BD (Breakspear et al., 2015; McLean et al., 2022) but also highlight a window for preventive strategies, where monitoring and potentially modulating these accelerated changes could mitigate subsequent clinical morbidity .

Also, we observed that adolescents at higher genetic risk for BD also showed increased subclinical psychiatric symptoms and cognitive-emotional alterations in their daily lives. These behavioral profiles align with prior evidence that neurodevelopmental disruptions in BD manifest in social, emotional, and cognitive dysfunction (Hafeman et al., 2017). For example, heightened impulsivity or emotional lability may affect academic performance and peer relationships, creating cumulative stressors that further exacerbate risk. Importantly, our network and cortical findings provide a structural and mechanistic context for these real-world functional challenges: adolescents with more globally integrated but less locally segregated networks may struggle to effectively filter and regulate emotional information, while cortical thinning in frontal areas could undermine executive control processes (Chang et al., 2012). Taken together, these results underscore the importance of integrating neuroimaging markers with behavioral assessments to fully grasp the multidimensional burden of BD risk, ultimately guiding tailored early interventions that address both neurobiological vulnerabilities and their day-to-day manifestations.

Finally, we observed that the expression patterns of three candidate genes for BD—PLEKHA2, ZSCAN31, and ANK3—showed a high degree of spatial overlap with regions of cortical structural abnormalities identified in adolescents with PRS score. This suggests that these genes may mediate neurodevelopmental deviations in BD by modulating cortical maturation. Among them, ANK3 is a well-established core risk gene for BD, known to be involved in postsynaptic membrane stability and the regulation of neuronal excitability. Its high expression in the prefrontal and temporal cortices closely corresponds with the regional cortical thinning observed in our study (Biernacka et al., 2019; Fiorentino et al., 2014; Lim et al., 2014; Roby, 2017). PLEKHA2, predominantly localized to the postsynaptic membrane, may influence synaptic signal integration and plasticity, thereby impacting prefrontal cortical development (Uhlén et al., 2015). Although ZSCAN31 has been less extensively studied, as a zinc finger transcription factor, it may play a critical role in cortical neuron migration and the formation of neural connectivity (Pardiñas et al., 2018; The GTEx Consortium, 2020). Pathway enrichment analysis further revealed that these genes are highly expressed in key neuronal structures involved in synaptic transmission and connectivity, such as the postsynaptic membrane and dendritic spines, underscoring their biological relevance in neural circuits that regulate emotion and cognition (Andoh & Koyama, 2021). Consistent with previous findings on ANK3, this study is the first to link the expression profiles of PLEKHA2 and ZSCAN31 to neuroanatomical abnormalities in adolescents, thereby expanding our understanding of the mechanisms through which genetic risk for BD may exert its influence during neurodevelopment. Collectively, these findings not only highlight the potential for BD genetic susceptibility to shape atypical cortical developmental trajectories during adolescence, but also offer novel molecular insights for early identification, risk prediction, and targeted intervention—advancing the field toward precision psychiatry.

## Limitation

Several limitations warrant consideration in this study. First, this study’s sample was obtained from a single center, potentially restricting the generalizability of the findings; hence, future research should incorporate multi-center data to capture broader demographic and clinical variability. Although the ABCD dataset is large, dividing participants into high- and low-PRS subgroups yielded relatively small sample sizes in each group, potentially diminishing statistical power. Second, while we collected longitudinal data, the current time points may be insufficient to fully characterize the developmental trajectory of BD risk. Future studies should include more frequent follow-up assessments during critical stages of adolescence to obtain richer longitudinal information. Third, the gene expression data utilized in this study were derived from a limited number of independent human samples, with inherent gender- and hemisphere-related sampling imbalances. Future investigations could benefit from expanding the microarray dataset through inclusion of additional samples from publicly available resources. Such expansion would enable more robust validation of our current findings regarding candidate genes while facilitating the identification of risk genes associated with BD. This enhanced dataset would particularly allow for systematic exploration of potential neuroimaging patterns with gene expression related to BD pathophysiology.

## Conclusion

Our findings demonstrate that genetic risk for BD significantly impacts adolescent brain development, resulting in distinct cortical and network-level alterations detectable through neuroimaging. These neurobiological deviations, coupled with behavioral manifestations, offer valuable biomarkers for identifying individuals at heightened risk. Integrating genetic, neuroimaging, and behavioral approaches provides a robust framework for early risk stratification and targeted preventive interventions, moving towards personalized precision psychiatry for BD.

## Method

### Adolescent Brain Cognitive Development (ABCD)

Leveraging comprehensive demographic, genetic, and neuroimaging data from 4,579 unrelated neurotypical children (119.37 ± 7.34 month; 2117 females / 2402 males) from the multi-site ABCD 2.0.1 dataset release across 22 sites, we aimed to examine how genetic susceptibility to BD may influence brain structure within a large pediatric cohort. Participants were systematically recruited through probability sampling of schools geographically proximate to each study site. Parental or guardian informed consent, along with child assent, was obtained prior to participation. Ethical approval for this investigation was provided by the Institutional Review Board at the University of California, San Diego. The substantial size and diversity of this dataset present an unparalleled opportunity to clarify genetic determinants underlying BD risk within a neurotypical pediatric population. The demographics were shown in **Table S1**.

### Clinical Assessment and Substance use

In the ABCD study dataset (Michelini et al., 2019), we used scales from the Kiddie Schedule for Affective Disorders and Schizophrenia (K-SADS, *mh_y_ksads_bip*) to measure the severity of adolescence BD. These items are employed to gather and record information on participants’ BD-related symptoms.

The datasets *su_y_alc_exp*, *su_y_can_exp*, and *su_y_nic_exp* pertain to substance use expectancies, where *alc* refers to alcohol, *nic* refers to nicotine, and *can* likely refers to cannabis. The positive expectancies reflect an individual’s inclination toward substance use. Substance use represents positive expectancies toward nicotine, alcohol, and cannabis use, indicating the extent to which individuals anticipate positive experiences from substance use. Higher substance use scores suggest a greater likelihood of future substance use. The details of those dataset could be seen in *Supplementary Materials*.

### MRI data analysis

Minimally preprocessed T1-weighted MRI scans obtained from the ABCD dataset underwent cortical reconstruction using FreeSurfer software (version 6.0.1), following standardized processing protocols (Fischl, 2012). The cerebral cortex was parcellated into 68 distinct cortical regions based on the Desikan-Killiany atlas. Inter-subject alignment was achieved via surface-based registration of the T1-weighted images. Mean cortical thickness, serving as a macrostructural MRI-derived phenotype, was extracted and standardized to facilitate subsequent analyses. Morphological data were harmonized across sites using ComBat (https://github.com/Jfortin1/ComBatHarmonization), a post-acquisition statistical method that corrects for site-related batch effects while preserving biological variability related to age, sex, and genetic risk (Fortin et al., 2017). Quality control follows the process of previous study (Hagler et al., 2019) .

### Genetic Data

Genotyping data were generated using the Affymetrix Axiom Smokescreen Array platform (Baurley et al., 2016; Uban et al., 2018), with quality control procedures for both samples and variants previously detailed [https://github.com/neurogenetics/GWAS-pipeline]. Samples were excluded based on genotype missingness >1%, heterozygosity beyond ±3 standard deviations, sex discordance, or relatedness (π > 0.125). We restricted our analysis to individuals of genetically inferred European ancestry to ensure compatibility with the populations used in the reference GWASs. Genotype imputation was conducted via the Michigan Imputation Server using the HRC 1.1 2016 reference panel and Eagle v2.3 for phasing.

### Polygenic risk score for bipolar disorder

Polygenic risk scores (PRS) were computed for each individual based on summary statistics from a GWAS of BD (Mullins et al., 2021). SNPs with an imputation quality score *(INFO) <*

0.3 or *MAF <* 0.01 were excluded, and duplicate SNPs were removed prior to score computation. PRSice-2 was utilized to generate individual PRS, selecting SNPs with *p* < 0.1, consistent with the previously reported optimal threshold for BD polygenic risk estimation in GWAS studies (Mullins et al., 2021). This methodological framework ensures robust genotypic data processing, facilitates high-resolution imputation, and enables the derivation of PRS to elucidate the genetic architecture underlying BD susceptibility.

### Normative model for cortical thickness

The normative models for each neuroimaging measure—cortical thickness derived from FreeSurfer were developed using the CentileBrain normative modeling framework (Ge et al., 2024). The model parameters in the normative framework (CentileBrain; https://centilebrain.org) were applied to each regional cortical thickness measure of the individuals with BD. This study constructed normative models of cortical thickness using multivariate fractional polynomial regression (MFPR) based on data from 37,407 healthy individuals (aged 3–90 years) across 87 datasets from Europe, Australia, the USA, South Africa, and East Asia. The MFPR-based models demonstrated robustness to multisite effects, ethnic diversity, and variations in image processing software versions (Ge et al., 2024). For each measure in each participant, we estimated the degree of normative deviation from the reference population mean as a deviation Z-score. A positive or negative Z-score indicated that the value of the corresponding morphometric measure was higher or lower, respectively, than the normative mean. Previous literature (Ge et al., 2024b; Lv et al., 2021; Verdi et al., 2023), we defined regional Z-scores as infranormal when below −1.96 or supranormal when above +1.96, corresponding to the 5th percentile and 95th percentile, respectively.

### Structural covariance and topographical property

Individual covariance networks were constructed by calculating product-moment correlation coefficients of cortical thickness across different brain regions, thereby capturing interregional structural characteristics (DuPre & Spreng, 2017). We derived two global metrics: (1) mean clustering coefficient, which quantifies the local interconnectedness of brain regions, and (2) mean path length, which reflects the average shortest path between any two regions in the network (Bullmore & Sporns, 2009; He et al., 2007). At the regional level, we employed a similar strategy to that used for the global metrics. Specifically, for each cortical and subcortical region in the thresholded structural covariance network, we computed normalized clustering coefficient, normalized path length, and their ratio. Based on effect size distributions, we generated topological feature maps to identify regions that deviate from a “small-world” organization (Larivière et al., 2022). For instance, regions displaying increases in both clustering coefficient and path length were characterized as “regularized,” whereas those exhibiting decreases in both metrics were considered “randomized.” This approach provides a more intuitive framework for assessing the structural properties and potential abnormalities of brain networks across multiple dimensions (Larivière et al., 2022).

### Longitudinal Changes in Brain Structure

Furthermore, we utilized longitudinal cortical thickness measurements and corresponding structural covariance network properties to further assess the rate of morphological brain changes between high- and low-PRS groups. Specifically, we defined the rate of brain structural change as the difference in cortical thickness and structural covariance network metrics—namely, clustering coefficient and shortest path length—between baseline and the two-year follow-up. Note that ΔCT was calculated as the cortical thickness measured at the two-year follow-up minus the baseline cortical thickness within each individual subject. Subsequently, we compared these structural change rates between the high-risk and low-risk groups, while controlling for potential confounding variables, including age, sex, and handedness.

### Association between PRS and Substance Use Exposure

We aim to compare differences in substance positive expectation between the high PRS group and the low PRS group. To assess group differences, we performed an *independent two-sample t-test*, controlling for age, sex, head motion and right-handedness as covariates.

### Cortical thickness predicts Substance Use Exposure

To further clarify the relationship between cortical thickness and biomarkers of Substance Use Exposure Expectation, we employed a multivariate machine learning model for predictive analysis of behavioral data. Specifically, we utilized a Support Vector Regression (SVR) modelvia Scikit-learn (Pedregosa et al., n.d.) to predict participants’ substance use exposure based on extracted cortical thickness features from specific brain regions. Initially, we divided the dataset (*N* = 474) into training and testing subsets using 10-fold cross-validation to mitigate model overfitting. For each cross-validation iteration, we trained the SVR model using default parameters and a linear kernel. Predictions from each fold were then recorded and combined. Prediction accuracy was assessed by calculating the Pearson correlation coefficient between the predicted and actual values. To evaluate the stability and robustness of our model, we repeated the 10-fold cross-validation procedure 1000 times. The predicted outcomes for each sample across all iterations were averaged to obtain the final prediction results, ensuring reliability and reproducibility of the model predictions.

### Genetic mechanism

To elucidate the genetic regulation underlying structural brain abnormalities, we deconvolved cell-type fractions from microarray data obtained from the Allen Human Brain Atlas (AHBA; http://human.brain-map.org/) (Shen et al., 2012). Integrating these transcriptomic profiles with high-resolution brain connectivity architecture, we aimed to investigate the molecular mechanisms influencing cortical thickness alterations. The microarray dataset included 3,702 spatially distinct samples derived from six post-mortem adult human brains (mean age = 42.5 ± 13.38 years; male / female = 5 / 1). Gene expression data were processed and parcellated into 68 cortical regions based on the left Desikan-Killiany (DK68) atlas using the Abagen toolbox (https://github.com/netneurolab/abagen) (Markello & Misic, 2021). Spatial correlations were computed between regional gene expression and cortical thickness patterns derived from neuroimaging data. Genes demonstrating significant associations were subsequently cross-referenced with those identified in a large-scale GWAS of BD (Mullins et al., 2021). Overlapping genes were selected for downstream analysis to elucidate convergent molecular signatures. We further investigated whether genes exhibiting transcriptional dysregulation in BD postmortem brain tissue (as reported in RNA-seq datasets, e.g., PSYCHENCODE) showed preferential expression in cortical regions implicated in cortical thickness abnormalities. To functionally annotate these candidate genes, we performed pathway enrichment analysis using Metascape (https://metascape.org), a platform that integrates over 40 curated biological databases (Zhou et al., 2019). Enrichment significance was assessed using a null-model-based approach, applying a false discovery rate (FDR) correction threshold of *q* = 0.05.

## DATA AVAILABILITY

The ABCD dataset could be accessed at https://nda.nih.gov/abcd. Allen Human Brain Atlas could be accessed at http://human.brain-map.org/, ENIGMA toolbox (https://enigma-toolbox.readthedocs.io/en/latest/pages.html).

## CODE AVAILABILITY

Code will be available on https://github.com/Laoma29/Publication_codes.

## ACKNOWLEDGMENTS

Xiaobo Liu and Jiadong Yan are supported by the China Scholarship Council. Bin Wan is supported by International Max Planck Research School on Neuroscience of Communication: Function, Structure, and Plasticity (IMPRS NeuroCom), Graduate Academy Leipzig, and Mitacs Globalink Research Award.

## ABCD Dataset

Specifically, the alcohol expectancy scale (su_y_alc_exp) includes items such as: “Drinking alcohol would make it easier to make friends,” “Drinking would help me relax,” “Drinking would make me feel happier,” “Drinking would give me more confidence,” and “Drinking would help me fit in better socially.” The nicotine expectancy scale (su_y_nic_exp) includes: “Smoking would help me relieve stress,” “Smoking would help me concentrate,” “Smoking would make me look cool,” “Smoking would make it easier to make friends,” and “Smoking would help me relax.” The cannabis expectancy scale (su_y_can_exp) comprises items such as: “Using marijuana would make me feel more relaxed,” “Marijuana would make me more creative,” “Using marijuana would help me fit in better socially,” “Marijuana would make me feel happier,” and “Using marijuana would help me escape from my problems.” Participants rated their level of agreement with each statement using a Likert-type scale (e.g., from “strongly disagree” to “strongly agree”).

**Table 1.** Demographic characteristics of the study population.

|  | Number | Age<br>(Month<br>) | Sex<br>(Female/Male) | Handedness<br>(Right/Mixed/Left) | su_y_alc_exp | su_y_can_exp | su_y_nic_exp |
| --- | --- | --- | --- | --- | --- | --- | --- |
| Total | 4519 | 119.37<br>±7.34 | 2117/2402 | 3588/323/608 | 6.05±2.69 | 4.47±2.25 | 6.68±2.81 |
| High PRS | 237 | 118.27<br>±7.20 | 101/137 | 186/16/36 | 7.23±2.88 | 7.51±2.81 | 4.81±2.65 |
| Low PRS | 237 | 120.54<br>±6.97 | 114/123 | 185/17/35 | 6.61±2.73 | 6.91±3.00 | 4.64±2.22 |

